# Modelling of filamentous phage-induced antibiotic tolerance of *P.aeruginosa*

**DOI:** 10.1101/2021.12.03.471163

**Authors:** Maria van Rossem, Sandra Wilks, Malgosia Kaczmarek, Patrick R. Secor, Giampaolo D’Alessandro

## Abstract

Filamentous molecules tend to spontaneously assemble into liquid crystalline droplets with a tactoid morphology in the environments with the high concentration on non-adsorbing molecules. Tactoids of filamentous Pf bacteriophage, such as those produced by *Pseudomonas aeruginosa*, have been linked with increased antibiotic tolerance. We modelled this system and show that tactoids, composed of filamentous Pf virions, can lead to antibiotic tolerance by acting as an adsorptive diffusion barrier. The continuum model, reminiscent of descriptions of reactive diffusion in porous media, has been solved numerically and good agreement was found with the analytical results, obtained using a homogenisation approach. We find that the formation of tactoids significantly increases antibiotic diffusion times leading to stronger antibiotic resistance.

## Introduction

Polymicrobial biofilms, where microcolonies of bacteria form structured communities surrounded by a self-produced extracellular polymeric substances (EPS), are increasingly linked to infections and to providing an environment that encourages bacterial survival and persistence. Bacterial survival in biofilms is supported by parameters such as lower metabolic rates, stress responses, decreased nutrient diffusion and low oxygen gradients.

Recent work [1–4] on structuring and organisation of such systems has found that biofilms can exhibit order comparable to liquid crystals. The formation of the liquid crystalline order was also detected in other biological materials including cell tissue [5–7] and cellulose [8–10]. While, in general, the mathematical modelling of biological liquid crystals is a well-explored topic [10], for biofilms the models are limited. As this a relatively new area, the mathematical approaches developed so far mainly concern global population dynamics [11] and empirical fitting models [12–17], with some recent, pioneering work also including cell-level simulations [3, 4]. To the best of our knowledge, no continuum models describing liquid crystalline order in biofilms exist.

The latest results by Secor et al. (2015) [1] have shown that, upon addition of non-adsorbing polymers such as DNA to the *Pseudomonas aeruginosa* filamentous phage (Pf4), a phase separation occurs, with phages forming a nematic liquid crystalline phase consisting of droplets with a rugby ball shape, called tactoids (Fig 1), surrounded by an isotropic phase predominantly consisting of the non-adsorbing polymers. Although the tactoid morphology has not been directly observed in biofilms, nematic structure has [1]. Due to their uniform properties such as length, diameter, and charge, filamentous phages (with or without non-adsorbing polymers) can provide an experimental model system to study the physics of liquid crystal formation and depletion attraction, which has been described previously [18–20].

**Fig 1.**
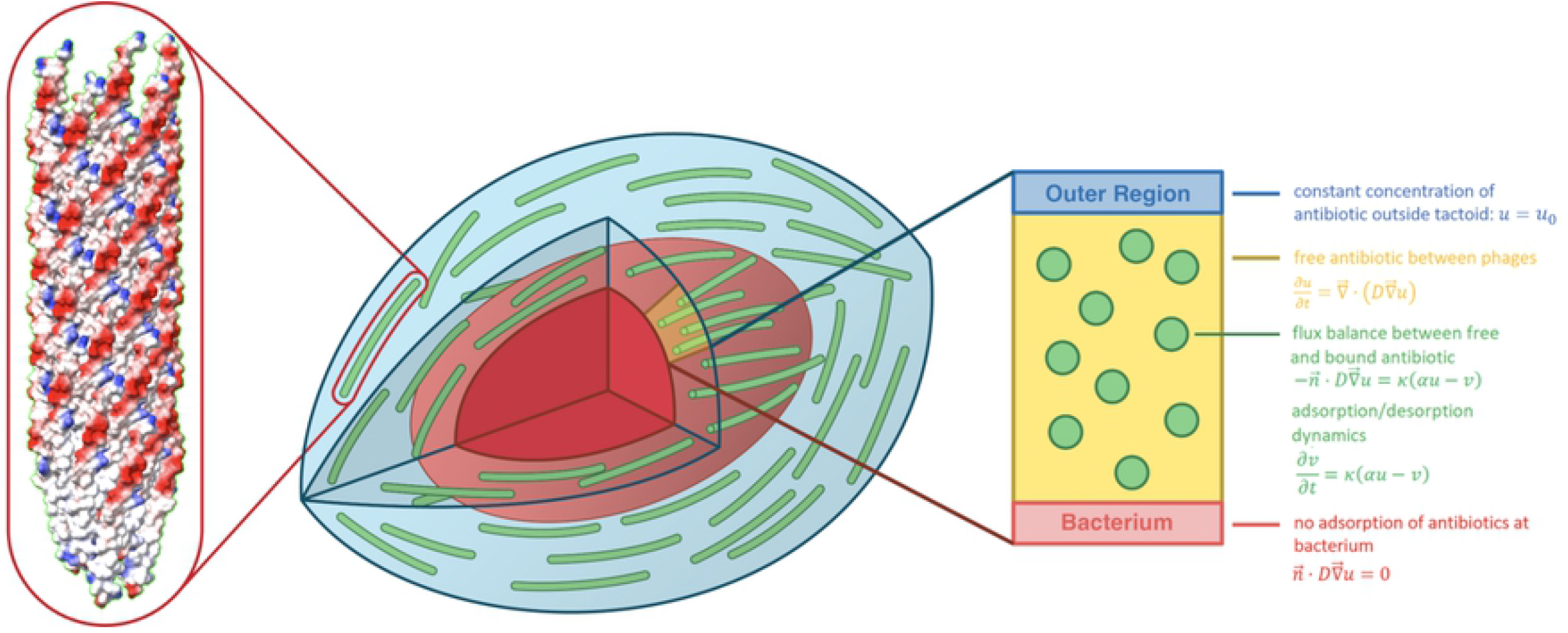
Schematic sketch of the tactoid model. The central image shows a schematic sketch of a tactoid (in blue), with phages (green) surrounding a bacterium (red). The structure of a phage is shown on the left, with colour indicating electric charge: red signifies negative charge, and blue positive. An example of a two-dimensional tactoid domain on which the model is solved is shaded in yellow. This domain is sketched on the right, with model equations and domain parameters indicated.

The Pf4 phage (genus Inovirus) is approximately 2 μm in length and 6 – 7 nm in diameter. It is a long, negatively charged filament with more uniformity than synthetic filamentous nanoparticles. *P. aeruginosa* itself is an important bacterial pathogen, causing a wide range of infections including wound and burn infections, pneumonia, urinary tract infections, and it is a dominant pathogen in cystic fibrosis airway infections. *P. aeruginosa* can readily form biofilms, which are often characterized by a highly mucoid EPS containing polysaccharides, proteins, lipids and DNA. Further research has shown that Pf virions encapsulate *P. aeruginosa* cells [2].

The production of Pf phages by *P. aeruginosa* is stimulated by high viscosity environments [21], anoxic conditions [22], and oxidative stress [23], conditions that mimic those found at infection sites. Pf bacteriophages are often found in cystic fibrosis sputum samples [24] and have a direct role in biofilm formation [25, 26]. Therefore, the behaviour of these phages in the presence of polymers (representing those which could be found in the EPS) offer a model to start understanding how Pf virions in environments that facilitate liquid crystal formation could impact tolerance and antibiotic diffusion.

There is growing concern over the increasing prevalence of antibiotic resistance and tolerance. Resistance requires genetic mutation, while tolerance relates to an increased capability to be able to withstand antibiotic exposure temporarily, for example through the effects of decreased diffusion caused by the EPS. In *P. aeruginosa* biofilms, liquid crystal formation by Pf virions leads to increased antibiotic tolerance [1]. The ability of Pf virions to encapsulate *P. aeruginosa* cells suggests that a possible cause of the increased tolerance is this protective phage barrier [2]. It has been proposed that binding of cationic antibiotics by anionic Pf virions (observed by Janmey (2014) [27] and Secor et al. (2015) [1]) in a liquid crystalline state plays a central role in mediating antibiotic tolerance [1]. On the other hand, Pf virions do not provide protection from uncharged antibiotics such as ciprofloxacin, which is not bound by Pf4 phage [28].

The main focus of our work is to determine whether tactoids can cause antibiotic tolerance by acting as an adsorptive diffusion barrier without any further biological mechanisms. We approach this problem from a physical and mathematical angle by modelling diffusion and adsorption of antibiotics in a tactoid. The model presented here describes the diffusion of antibiotics using continuum equations for diffusion in a domain perforated by a regular lattice of filamentous phages, with adsorption at the phage boundary; this domain represents the tactoid (see Fig 1). It is solved numerically and compared against an analytical approximation, which is derived using the principle of homogenisation [29]. In homogenisation, scale separation and local periodicity at a microscopic scale lead to effective macroscopic equations. The technique is well developed in application to porous media [29–36], where it leads to estimating effective diffusion coefficients and time scales. For instance, homogenisation has been used to model adsorption-induced blockage of porous filters [37] and obstructed diffusion in polymer solutions [38]. Its application to adsorption and diffusion of antibiotics in the phage liquid crystalline phase allows us to estimate analytically the effective antibiotic diffusion time and, hence, to estimate the tactoid barrier efficiency.

Experiments have shown that adsorption leads to agglomeration of cationic antibiotics in tactoids [1]. Since this coincides with tactoids forming an effective barrier against such type of antibiotics, we predict that antibiotics will agglomerate in the outer tactoid layer before diffusive equilibrium sets in.

## Methods

Diffusion of antibiotics through a single tactoid encapsulating a bacterium was captured in a mathematical model and expressed through a system of dynamical equations. These equations were solved numerically in Comsol, to assess the influence of the tactoid on the diffusion time. The mathematical model, and its numerical implementation, are discussed in the following paragraphs.

### Description of the model

We describe the diffusion and adsorption of antibiotics by a system of equations analogous to diffusion in porous media, as described by Allaire et al. (2010) [29]:

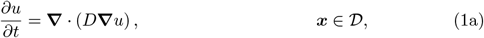

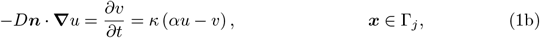

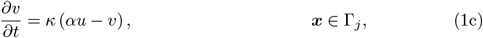

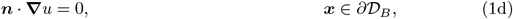

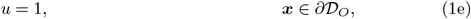

where *u* denotes the volume concentration of free antibiotics (in units of m^-3^), *v* is the surface concentration of adsorbed antibiotics (in units of m^-2^), *D* is the diffusion coefficient in m^2^/s, *α* is the equilibrium adsorption coefficient in m, which quantifies the degree of adsorption, *κ* is the adsorption rate in 1/s, and ***n*** is the unit normal pointing into the phages. Eq (1a) is a diffusion equation for the antibiotics in the tactoid, where *𝒟* is the tactoid volume not occupied by phages. Eqs (1b) and (1c) describe, respectively, the antibiotic flux and adsorption at the phage boundaries Γ_*j*_ (where the index *j* denotes the *j*-th phage), and Eq (1d) is a no-flux boundary condition at the bacterium boundary *∂𝒟*_*B*_, and Eq (1e) is the Dirichlet boundary condition at the outer tactoid boundary *∂𝒟*_*O*_. If needed, the no-flux boundary condition can be replaced by other conditions without affecting the mathematical analysis of the model. An illustration of the tactoid system, with the system of equations Eq (1) indicated, is shown in Fig 1. The system of equations Eq (1) is solved in Comsol. Since the aim of the model is to reproduce the experimental observations by Secor et al. (2015) [1], with some additional results by Tarafder et al. (2020) [2], the choices of not only the model, but also the values of its parameters (outlined in S2 Appendix) are based on the experimental work presented in these papers.

### Numerical implementation

Modelling an entire tactoid in three dimensions is computationally expensive. However, it is not necessary to do so, because a simpler modelling domain naturally follows from the geometry of the tactoid. The phages are very long, aligned with the bacterium, and have a diameter much smaller than that of the bacterium. The first two properties imply that antibiotic diffusion is only significant across the phages and not along them, thus reducing the problem from three to two dimensions. The last property allows us the neglect the curvature of the bacterium and to consider only a thin section of the tactoid (shaded in yellow in Fig 1) with periodic horizontal boundary conditions, so that the tactoid slice is effectively one layer in a vertical stack of identical slices. In summary, we can model the tactoid as a rectangular domain of length equal to the tactoid thickness, as shown in Fig 2. We assume that the phages form a regular lattice in the tactoid and choose the height of the rectangle to be the period of this lattice. This is a standard assumption in models of diffusion in porous media, made for numerical convenience [39, 40]. The antibiotics come in from the right, with the condition that initially no antibiotics are present inside the domain. The antibiotics diffuse to the left until a diffusive equilibrium is reached, i.e. a homogeneous distribution of antibiotics throughout the domain. We assume adsorption has a negligible effect on the concentration of antibiotics outside the tactoid, and set a constant concentration of antibiotics on the outer right boundary of our domain.

**Fig 2.**
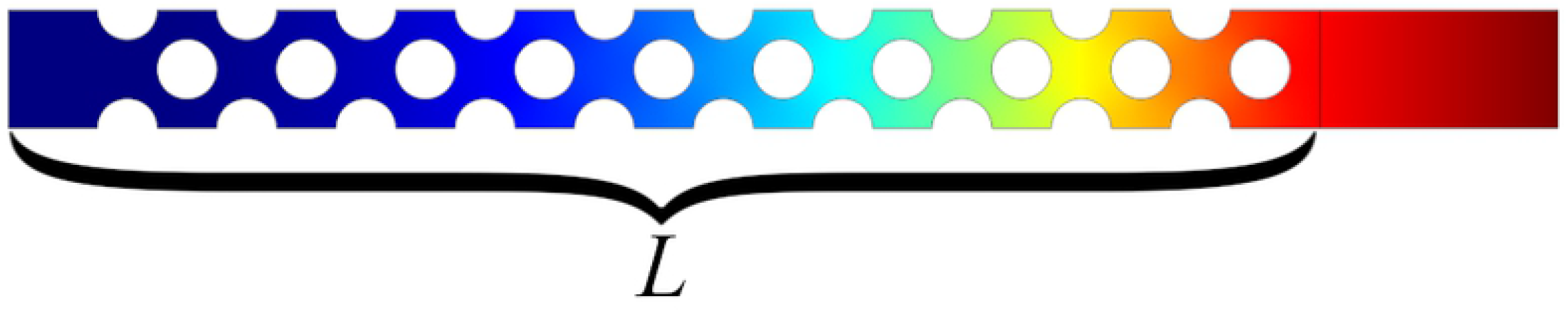
An example of the Comsol model geometry. The circles represent phages and the colour scale indicates antibiotic concentration (with high to low concentration represented by the colours from red to blue). The phages form a lattice of repeating unit cells. The tactoid width *L* is indicated.

## Results

The diffusion of the antibiotic tobramycin through an adsorptive tactoid layer was modelled, and the timescale at which antibiotics diffuse through to the bacterium was estimated as *t*_90_, the time at which the free antibiotic concentration at the bacterium boundary is 90% of the equilibrium concentration inside the domain. The modelling parameters, summarised in Table 1, were estimated from the experimental conditions in [1] and [2], and the equilibration times were compared with their experimental results. A full account of the parameter estimation is given in S2 Appendix.

**Table 1.** Summary of parameter values.

| Description | Symbol | Value |
| --- | --- | --- |
| nematic equilibrium adsorption coefficient | $\alpha_N$ | 2.2 $\mu\text{m}$ |
| isotropic equilibrium adsorption coefficient | $\alpha_I$ | 0.4 $\mu\text{m}$ |
| non-adsorptive equilibrium adsorption coefficient | $\alpha_0$ | 0 $\mu\text{m}$ |
| tactoid width | $L$ | 1 $\mu\text{m}$ |
| phage radius | $a$ | 3 nm |
| unit cell size | $L_c$ | 12 nm |
| diffusion coefficient | $D$ | 15 $\mu\text{m}^2/\text{s}$ |
| adsorption rate | $\kappa$ | $1.7 \times 10^6 \text{ s}^{-1}$ |

Since experiments show that the adsorptive capacity of the phages is different in the isotropic and nematic phases [1], the equilibration time was determined for multiple values of the equilibrium adsorption coefficient *α* (*α*_*I*_, *α*_*N*_ and *α*_0_ in Table 1), corresponding to isotropic, nematic and zero adsorption respectively. In the last case the phages merely form a physical barrier. The results are shown in Fig 3 and summarised in Table 2. The equilibration time without adsorption (*α* = 0) is close to the macroscopic diffusion time for tobramycin over the width of a tactoid in the absence of phages,

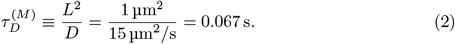

The numerical diffusion times (see Table 2) can be compared to the analytical effective diffusion times obtained by homogenisation, through the diffusion coefficients of the effective homogenised equations. The applicability of this method is demonstrated in Fig 3, which shows the increase of the antibiotic concentration at the bacterium boundary over time, relative to the equilibrium concentration. The concentration grows until it reaches this equilibrium value; the equilibration is significantly faster for weaker adsorption. The results of the microscopic and homogenised model are compared, showing good agreement between the two. The diffusion time, as calculated from the homogenised model, also agrees well with the numerical equilibration time, see Table 2.

**Fig 3.**
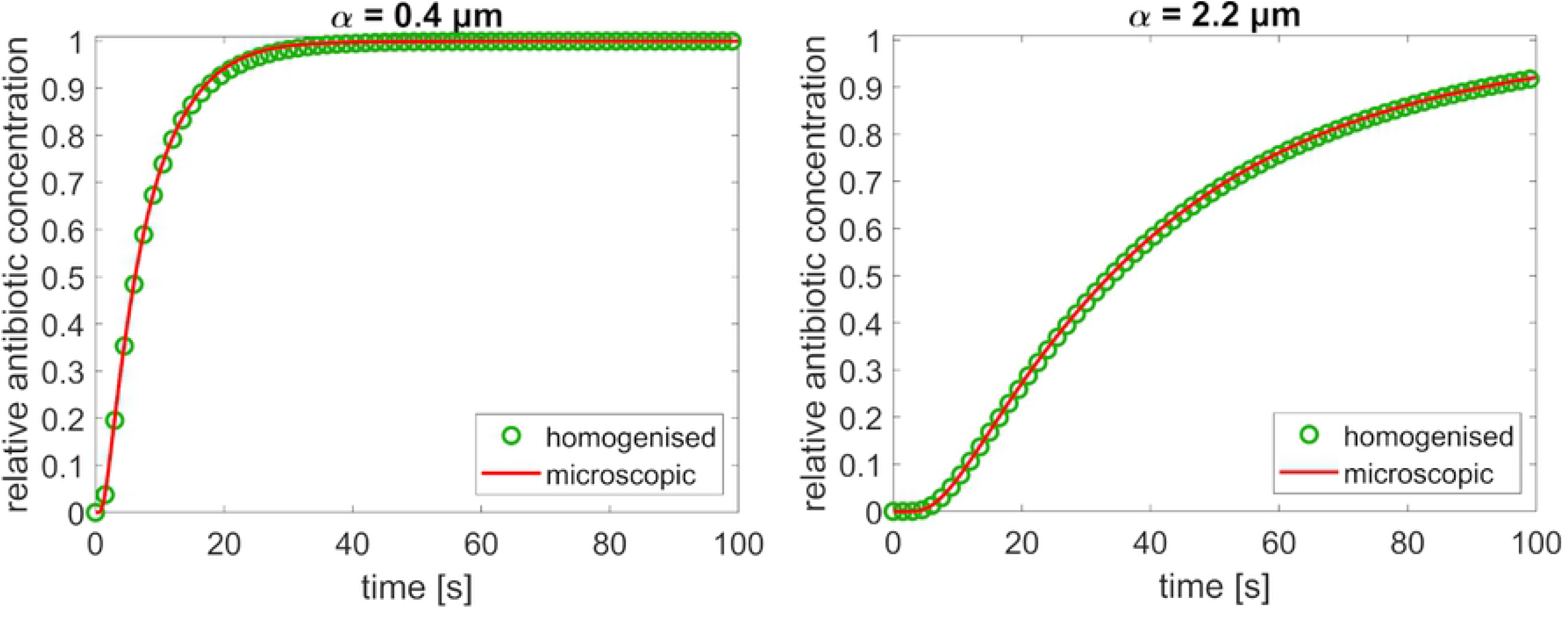
The antibiotic concentration at the bacterium boundary over time, as a fraction of the equilibrium concentration, with a comparison of the microscopic and homogenised model. The results are shown for two degrees of adsorption, quantified by the binding equilibrium constants *α*. The number of phage layers in both graphs is N=83, corresponding to a tactoid thickness of 1 μm and an interphage distance of 6 nm.

**Table 2.** Equilibration time for various values of the binding equilibrium constant *α*.

| $\alpha$ ($\mu\text{m}$) | $t_{90}$ (s) | $\tau_D^H$ (s) |
| --- | --- | --- |
| 0 | 0.112 | - |
| 0.4 | 16.6 | 16 |
| 2.2 | 93 | 89 |

To assess the agglomeration of antibiotics, the distribution of antibiotics across the tactoid was modelled and plotted after 10 s, before diffusive equilibrium has set in, and after 100 s, when the system has (almost completely) reached equilibrium. These results are shown in Fig 4 and indicate that antibiotics agglomerate in the tactoid: at equilibrium, the concentration in the tactoid is higher than outside of it. Furthermore, antibiotics agglomerate in the outer tactoid layer before equilibrium has set in; this effect is transient and more pronounced for stronger binding equilibrium coefficient.

**Fig 4.**
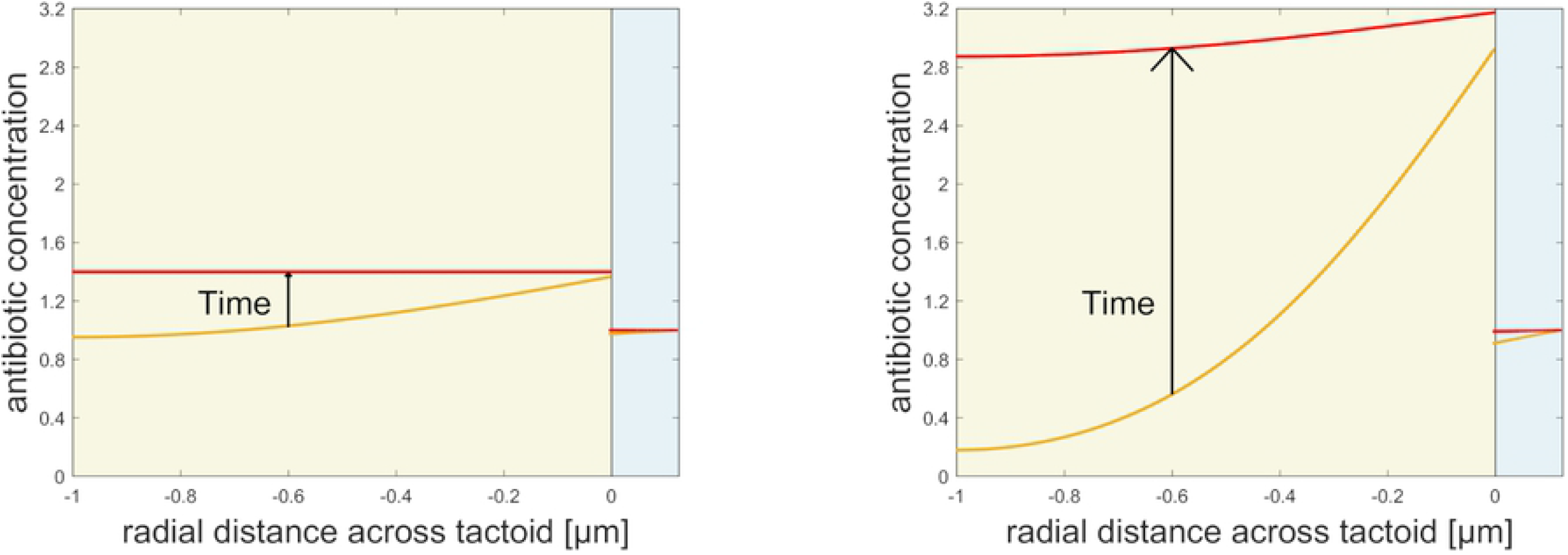
Distribution of antibiotics (both free and adsorbed) across the tactoid. Left and right figures show different equilibrium adsorption coefficients: left is *α* = 0.4 μm (isotropic), right is *α* = 2.2 μm (nematic). Each figure shows the antibiotic concentration distribution after 10 s (orange line), and 100 s (red line). The time progression is indicated by an arrow. The distance covered ranges from the bacterium boundary at *−* 1 μm, to just beyond the outer tactoid boundary at 0 μm; the tactoid domain is indicated by a yellow background and the outer domain is blue. The discontinuity in the antibiotic concentration of the outer tactoid boundary is due to the lack of bound antibiotics outside the tactoid.

Values are obtained from numerical simulation of the system of equations Eq (1) (*t*_90_) and from the homogenised model 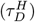.

## Discussion

The model outlined in this paper is an extremely powerful tool to link experiments that measure phage parameters, e.g. adsorption, to antibiotic resistance. The case discussed in the previous section is just an example, based on parameter values estimated from the results reported in [1] and [2]. However, the power of the model lies in its ability to link changes in the experimental conditions to antibiotic resistance. For example, increasing the ion concentration in the solution creates more compact tactoids; a larger affinity between phages and antibiotic can increase the amount of adsorbed antibiotic. Both effects can lead to an increased antibiotic tolerance, but our model allows us to quanitfy by how much. The increase can be infinite for the former effect, but is only linear for the latter. In this section we analyse in turns the effect of the various model parameters on the antibiotic diffusion time, using it as a proxy for antibiotic resistance. All the numerical results presented are obtained using the numerical integration of Eq (1), but we use the homogenised model to guide us in their interpretation.

We start by analysing the effect of tactoid thickness *L*. The homogenised model outline in S1 Appendix, Eq (3), is equivalent to diffusion in a domain without phages with an effective diffusion coefficient *D*^(eff)^. Therefore the diffusion time is given by equation (2) with the microscopic diffusion coefficient *D* replaced by *D*^(eff)^. From this equation we see immediately that the diffusion time increases quadratically with the domain size, a fact that is confirmed by the numerical integration of Eq (1), see Fig 5.

**Fig 5.**
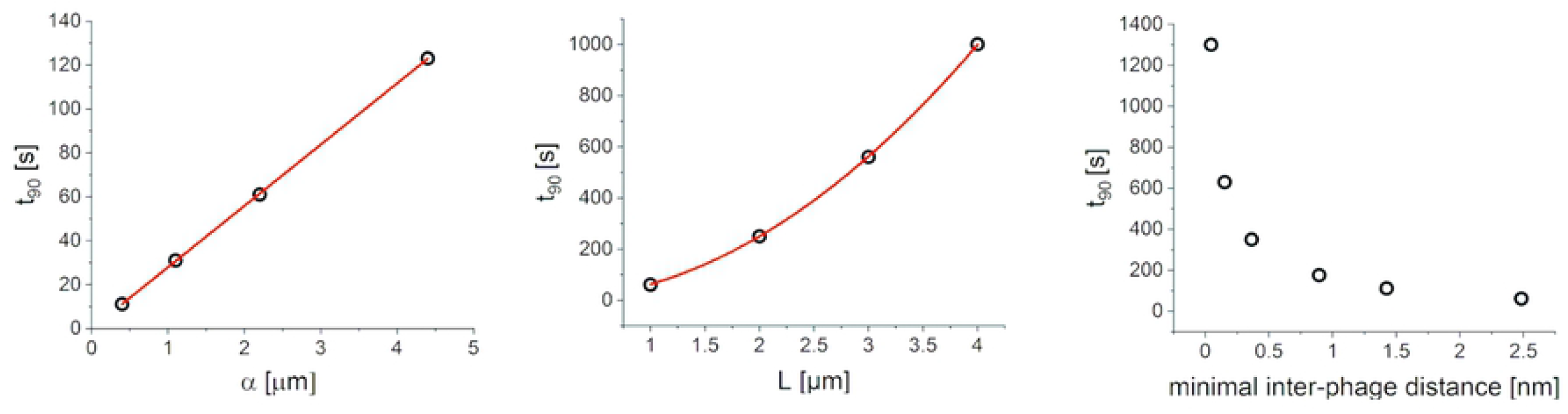
Dependence of equilibration time on model parameters. The dependence is shown for (a) the binding equilibrium constant *α*, with a linear fit indicated by the red line (b) the tactoid width, with the red line showing a quadratic fit, and (c) the phage packing density.

The effect of the equilibrium adsorption coefficient *α* can also be analysed through the effective diffusion time 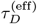 of the homogenised model. In the limit of small *α*^*−*1^, 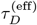 scales linearly with *α* (see S1 Appendix for more details). This linear relationship is again confirmed numerically, as indicated in Fig 5.

The equilibration time shows a rapid increase with packing density, as shown in Fig 5. This agrees with intuition; as the phages approach the most efficient packing, the minimal interphage distance approaches zero and no gaps are left for the antibiotics to diffuse through. Hence, in the limit of zero interphage distance, equilibration time should diverge.

Finally, the results do not depend on the adsorption rate *κ*, as long as it is fast compared to the diffusion rate. Indeed, when varying *κ* from 100 */*s to 10^6^*/*s, the equilibration time does not change at all.

The numerical analysis of parameter effects summarised in Fig 5 was repeated for times at which the concentration at the bacterium is 10% of the average instead of 90%, to check that the results are not an artefact at 90%. The results agree well, as shown in Fig S2 Fig of the supporting material.

Having analysed the effect of the parameters, we can better assess the calculated diffusion times. The results in Table 2 show that encapsulation of bacteria by tactoids composed of Pf virions strongly increases antibiotic diffusion time, indicating that tactoids serve as an effective diffusion barrier. However, the resulting diffusion times, of the order of tens of seconds to minutes, are not sufficiently long to fully explain the observed antibiotic tolerance over timescales of tens of minutes to a few hours [1]. On the other hand, Table 2 shows that adsorption is essential to the efficacy of the tactoids as a diffusion barrier, and this agrees with the observed correlation between adsorption and antibiotic resistance [1, 2, 28]. Furthermore, the results in Fig 4 are consistent with the observation that antibiotics agglomerate in tactoids [1, 2]. These results reinforce the observation that the barrier effect of the tactoid directly impacts antibiotic tolerance [2], and support the hypothesis that diffusion inhibition and adsorption are sufficient for the barrier to be effective. Moreover, they provide a mechanism for predicting the influence and effect of this tactoid barrier.

The quantitative difference between modelled and observed tolerance timescales may be explained by underestimation of parameter values, the influence of adsorption on global antibiotic concentration, and the combined effect of tactoid encapsulation with other tolerance mechanisms. We will discuss these in turn.

The analysis of parameter effects earlier in this section shows that, while *κ* plays little role in antibiotic resistance, underestimating *α*, the packing density, or the tactoid thickness has a significant effect. We outline in S2 Appendix the procedure we followed to estimate the parameter values used to obtain the data in Table 2. In all cases more targeted experiment will be needed to constrain the parameters further. For example, we may have underestimated the values of *α*, since these were derived from measurements at a macroscopic scale; further experiments could elucidate how well these results carry over to the scale of a tactoid. Furthermore, the authors are currently developing a model to investigate why phages in the nematic tactoid phase have a stronger binding capacity *α* than in the isotropic phase, which was observed by Secor et al. (2015) [1]. However, the effect of *α* is relatively modest (see Fig 5). The remaining two parameters can potentially have a much larger impact. The phage packing density is dependent on the ionic strength of the environment (due to the anionic phage charge) and the molecular weight of the polymers serving as depletion agents; this yields an opportunity for future experiments to test the effect of packing density on antibiotic tolerance.

It is also possible that adsorption lowers the global antibiotic concentration enough for it to become non-lethal. However, to what extent this concentration is lowered depends on the number of tactoids present and is therefore not straightforward to estimate. The antibiotic concentration is lowered further for two reasons: in the first place, most biological polymers are anionic and can bind cationic drugs, which means that the EPS would adsorb part of the antibiotics *in vivo*. Secondly, antibiotics have half-lives, especially when enzymes are present to cause degradation, or when they are metabolized by the host.

Finally, the observed increase in antibiotic tolerance [1] may be a combination of various mechanisms. The results in Table 2 show that tactoids cause a slow arrival of low concentrations of antibiotics, which may trigger other tolerance and resistance mechanisms such as efflux, stress responses such as the SOS response to DNA damage, etc. that stop bacterial growth, protecting them from antibiotics that kill replicating bacteria. The slow arrival of antibiotics can also afford the bacteria temporary protection while these other mechanisms are not yet active. Furthermore, one should consider that in biofilms, not only the phages, but also the bacteria will aggregate, and microcolonies will form [41]. This means that the bacteria in the inner layers of the biofilm have more protection against antibiotics than one encapsulating tactoid can afford; the effect of tactoids around surrounding bacteria may be cumulative.

Possible interpretations of the aforementioned results hinge on the effect of liquid crystal formation on the development of antibiotic resistance of *P. aeruginosa*. Experiments by Burgener et al. (2019) show a correlation between antibiotic resistance and the presence of Pf phages in clinical samples, indicating that such an effect exists [28]. It would be of interest to verify whether it is the slow diffusion of antibiotics through tactoids (causing gradual exposure of bacteria to antibiotics) which induces this antibiotic resistance.

Furthermore, the model presented in this paper can potentially lead towards a further understanding of the complex role of the biofilm EPS matrix. In applying our results to *in vivo* biofilms, it is important to consider the experimental system on which our model has been based; a mix of Pf4 phages and non-adsorbing polymers which form tactoids have not been observed directly in biofilms. However, the biofilm EPS is likely to contain high amounts of similar non-adsorbing polymeric substances, and indeed there are additional host polymers in the system such as mucins or hyaluronan; therefore, we can assume that the entire biofilm matrix is a nematic liquid crystal (as suggested by Secor et al. (2015) [1]). Hence, the experimental system modelled in this paper offers a simplified model that can then be used to model the complex interactions between phage, bacteria, and polymers in polymicrobial biofilms where the role of the EPS matrix is only beginning to be understood.

In summary, our results indicate that tactoids can form a significant barrier against antibiotics, merely through adsorption and inhibition of diffusion. This result reproduces the trend observed experimentally and provides important insight into the physical mechanisms. In the next step of our work, we will consider the microscopic tactoid structure in a route towards more comprehensive, mathematical description of liquid crystalline biofilms.

## Conclusion

The model presented in this paper provides a clear quantitative link between geometrical and biochemical parameters and the antibiotic diffusion time through a liquid crystalline tactoid. It shows that tactoids can strongly increase the time it takes for antibiotics to reach encapsulated bacteria, by forming an efficient and antibiotic-specific adsorbing diffusion barrier. The fact that, without adsorption, the effect of the tactoids is negligible agrees with experimental observations of the lack of correlation between liquid crystal formation and the efficacy of ciprofloxacin, and previous conjectures [1, 28], as do the results on the agglomeration of antibiotics in tactoids [1]. The modelled increase in equilibration time is not sufficiently high to fully explain observed tolerance timescales [1]. This could be due to the underestimating of parameters or the neglected effect of adsorption on antibiotic concentration outside tactoids. However, our results are consistent with the previous work [1, 2] and indicate that encapsulation by tactoids acts as a barrier and may play a significant role in antibiotic tolerance. The results presented here offer a step towards better understanding of antibiotic tolerance in biofilms *in vivo*.

## Supporting information

**Fig S1.**
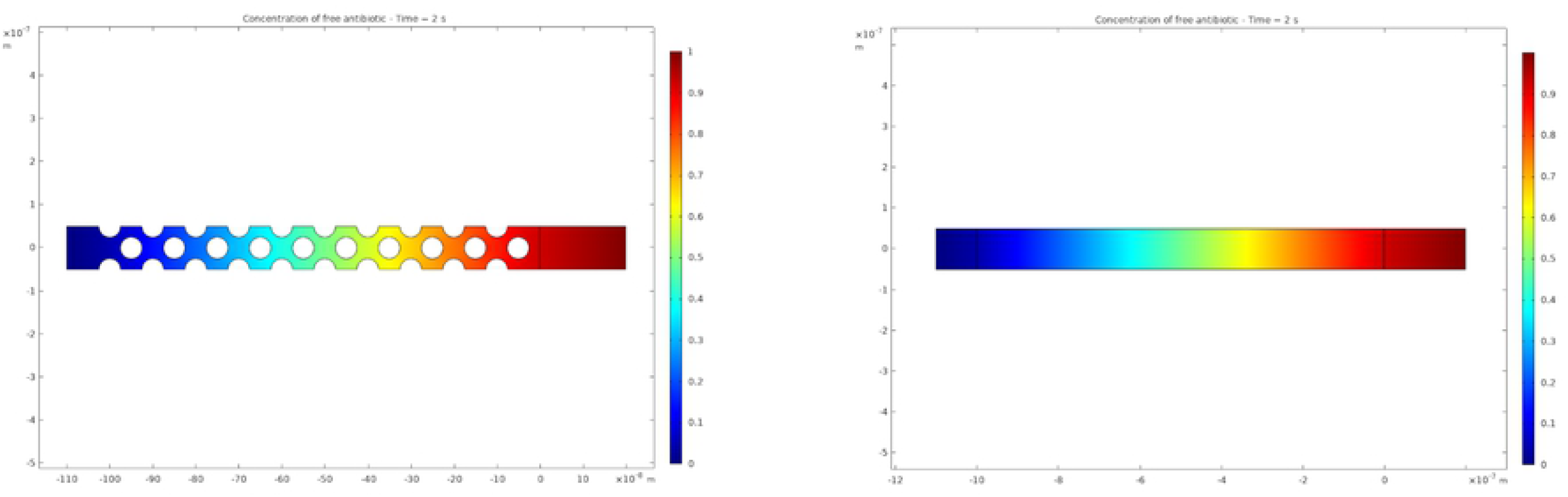
An illustrative comparison of the microscopic Comsol model geometry and the homogenised geometry without microscopic structure. The microscopic model is shown on the left; the homogenised model is shown on the right. The colour scale indicates free antibiotic concentration, circles represent phages.

**Fig S2.**
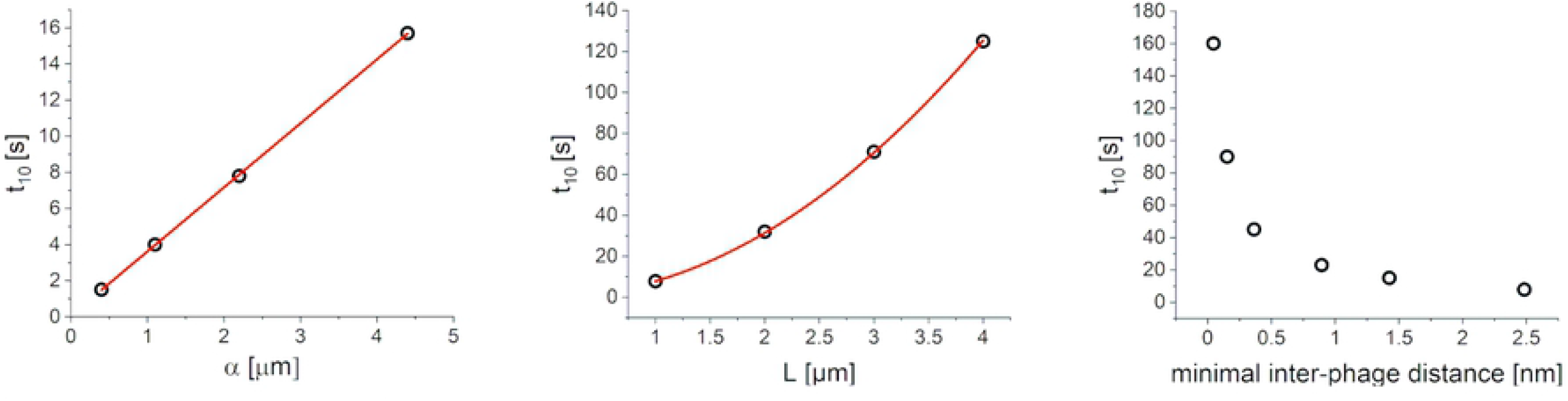
Dependence of *t*_10_ on model parameters. the dependence is shown for the binding equilibrium constant *α*, with a linear fit indicated by the red line (b) the tactoid width, with the red line showing a quadratic fit, and (c) the phage packing density.

## S1 Appendix Homogenisation

An analytically solvable effective model was developed using homogenisation. This is a mathematical technique used to describe systems in which scale separation occurs, i.e. where the physics at a macroscopic scale and a much smaller microscopic scale can be disentangled. Due to the microscopic structure, the system generally cannot be solved analytically, and numerical modelling is computationally expensive. However, in certain cases one can describe such a system system with effective equations on the macroscopic scale. If the microscopic details of the system are regular, their effect is averaged out and what is left is a much simpler macroscopic structure, governed by effective equations. These can often be solved analytically and give the same results as the microscopic model, in the limit that the microscopic scale becomes infinitesimal.

Homogenisation was applied to the microscopic model, following the work on diffusion in porous media by Allaire et al. (2010) [29]. The liquid crystalline alignment of phages leads to a regular structure which can be considered a lattice of identical unit cells: the microscopic structure to be averaged out. This process is illustrated in Fig S1 Fig. Homogenisation leads to the following effective equation:

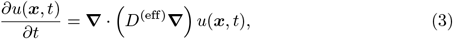

where *D*^(eff)^ is the effective diffusion coefficient, given by:

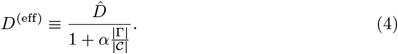

Here 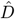 is a rescaled diffusion coefficient that takes into account of the physical barrier on diffusion caused by phages. The denominator, instead, represents the effect of adsorption. |*𝒞*| and |Γ| are the free volume between phages and phage surface in a unit cell, respectively.

This expression can be simplified in the limit of strong adsorption, i.e. (*α*|Γ|*/*|*𝒞*|) ≫ 1, to

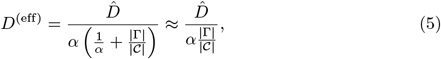

thus highlighting that the effective diffusion coefficient is inversely proportional to the equilibrium adsorption coefficient.

## S2 Appendix Parameter estimates

Model parameter values were set as follows. The tactoid width *L* was estimated from images by Secor et al. (2015) [1] and taken to be *L* = 1 μm. The phage radius *a* is 3 to 3.5 nm [1]; we use a conservative estimate of 3 nm. We set the unit cell size to be 4 times the phage radius; this value is estimated from Cryo-ET images by Tarafder et al. (2020), showing the configuration of phages inside a tactoid [2].

We take the diffusing antibiotics to be tobramycin, for which the greatest amount of data is available; its diffusion coefficient in an aqueous medium is *D* = 15 μm^2^*/*s.

The experiments by Secor et al. (2015) indicate an adsorption of about 30% of the antibiotics by phages in isotropic phase [1], which means *α* = 0.4 μm. When the phages are nematic, about 70% of antibiotics are adsorbed, which gives *α* = 2.2 μm.

Since we set *a* = 3 nm and *D* = 15 μm^2^*/*s we have a microscopic diffusion time

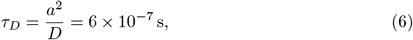

which, assuming that *τ*_*D*_ and *τ*_*κ*_ are of the same order, means *κ* ≃1*/τ*_*D*_ = 1.7 × 10^6^ s^−1^. Since *κ* is the adsorption rate, its value has no influence on the diffusion time, as long as *κ* ≫ 1s^−1^. This is confirmed by the fact that *κ* does not appear in equation (3). Hence, any large value for *κ* should give identical results.

A total antibiotic concentration of 200 μg*/*ml is used in the experiments of Secor et al. (2015) [1] which are comparable to the model presented in this paper. However, since only the relative antibiotic concentration is measured with respect to the average, *u* and *v* can be kept dimensionless. Furthermore, we assume that antibiotic adsorption has a negligible effect on the antibiotic concentration in the extracellular matrix surrounding the tactoid. Hence, we take the free antibiotic concentration at the outer tactoid edge to be *u* = 1.

## Acknowledgments

We want to thank Tim Sluckin for many insightful discussions

